# Patterns of genomic variation reveal a single evolutionary origin of the wild allotetraploid *Mimulus sookensis*

**DOI:** 10.1101/2023.09.14.557766

**Authors:** Makenzie R. Whitener, Hayley Mangelson, Andrea L. Sweigart

## Abstract

Polyploidy occurs across the tree of life and is especially common in plants. Because newly formed cytotypes are often incompatible with their progenitors, polyploidy is also said to trigger “instantaneous” speciation. If a polyploid can self-fertilize or reproduce asexually, it is even possible for one individual to produce an entirely new lineage. How often this extreme scenario occurs is unclear, with most studies of wild polyploids reporting multiple origins. Here, we explore the evolutionary history of the wild allotetraploid *Mimulus sookensis,* which was formed through hybridization between self-compatible, diploid species in the *Mimulus guttatus* complex. We generate a chromosome-scale reference assembly for *M. sookensis* and define its distinct subgenomes. Despite previous reports suggesting multiple origins of this highly selfing polyploid, we discover patterns of population genomic variation that provide unambiguous support for a single origin, which we estimate occurred ∼71,000 years ago. One *M. sookensis* subgenome is clearly derived from the selfer *M. nasutus*, which, based on organellar variation, also appears to be the maternal progenitor. The ancestor of the other subgenome is less certain, but it shares variation with both *M. decorus* and *M. guttatus*, two outcrossing diploids that overlap broadly with *M. sookensis*. Whatever its precise ancestry, this study establishes *M. sookensis* as an example of instantaneous speciation, likely facilitated by the polyploid’s predisposition to self-fertilize. With a reference genome for *M. sookensis* now available and its origin clarified, this wild tetraploid is poised to become a model for understanding the genetic and evolutionary mechanisms of polyploid persistence.

## Introduction

Polyploids are found across the tree of life and in nearly all extant flowering plant families (Jiao et al., 2011). This near ubiquity in angiosperms is likely due in part to the high incidence of polyploid formation (Barker et al., 2016; Herben et al., 2016; Kreiner et al., 2017; Ramsey, 2007), which is on the order of the mutation rate for autopolyploids (Ramsey & Schemske, 1998)but it might also be due to immediate and/or longer-term selective advantages of whole genome duplication (Comai, 2005). Nevertheless, the initial rarity of a newly formed polyploid (i.e., minority cytotype exclusion, Levin, 1975), combined with potentially strong reproductive isolation with its nearby progenitor species, presents a major challenge to establishment (Burton & Husband, 2000). One way to overcome this hurdle is self-fertilization (Rodriguez, 1996) and, indeed, polyploids are more likely to self than their diploid relatives (Barringer, 2007). In the extreme case of a single newly formed polyploid individual, its swift extinction is assured unless it can self-fertilize or backcross via a triploid bridge (Coyne & Orr, 2004; Ramsey & Schemske, 1998) – or unless there are other polyploid relatives nearby. Given this major constraint, it is perhaps not surprising that many polyploid species seem to have multiple evolutionary origins (reviewed in Soltis et al., 2014 and Soltis & Soltis, 1999). Even in self-fertilizing polyploids, which theoretically *can* evolve from a single individual, the high rate of unreduced gamete production (especially in F1 hybrids leading to allopolyploids, Ramsey & Schemske, 1998) might often facilitate multiple evolutionary origins.

After decades of studies reporting evidence of multiple evolutionary origins in diverse polyploid species (Abbott & Lowe, 2004; Allender & King, 2010; Doyle et al., 1990; Hegarty et al., 2012; Mavrodiev et al., 2015; Meimberg et al., 2009; Novikova et al., 2017; Segraves et al., 1999; Sigel et al., 2014; Vallejo-Marín et al., 2015; Wolfe et al., 2023; Zou et al., 2015), we might reasonably ask whether single-origin polyploidy is so rare as to be of little evolutionary importance. Still, there are notable exceptions to the multiple-origin rule including wild and cultivated species of peanut (*Arachis monticola* and *A. hypogaea*, Bertioli et al., 2019), *Coffea arabica* (Scalabrin et al., 2020), and the invasive allododecaploid *Spartina anglica* (Ainouche et al., 2004). It is also worth noting that most of what we know about the evolutionary origins of *wild* polyploid species comes from early molecular studies with a limited number of markers (Doyle et al., 1990; Hegarty et al., 2012; Mavrodiev et al., 2015; Segraves et al., 1999; Slotte et al., 2006; Soltis & Soltis, 1999; Soltis & Soltis, 1991). In the only wild polyploid systems with abundant population genomic data (species in *Arabidopsis* and *Capsella*), the question of evolutionary origins has been challenging to resolve. For example, in *C. bursa-pastoris*, support has been equivocal for multiple origins vs. a single origin with admixture from diploid relatives (Douglas et al., 2015; Kryvokhyzha et al., 2019), though a recent discovery of an early homeologous exchange event shared species-wide might provide evidence for the latter (Penin et al., 2023). In the allotetraploid *A. suecica*, early evidence from microsatellite markers had suggested a unique origin (Jakobsson et al., 2006), but more recent, whole-genome population resequencing revealed extensive shared polymorphism with diploid progenitors, suggesting multiple founding individuals (Novikova et al., 2017).

Given the technical challenges of phasing duplicated genomes, a reference assembly is an indispensable tool for gaining insight into the evolutionary origins of a polyploid species. However, it has been only recently that advances in long-read sequencing and chromatin confirmation capture have enabled assembly of complex and repetitive polyploid genomes (Michael & VanBuren, 2020). In polyploid crops, which include many species of major economic importance, the last few years have seen a burst of genome assembly (Bertioli et al., 2019; Borrill et al., 2015; Chen et al., 2020; Edger et al., 2019; Kamal et al., 2022; Scalabrin et al., 2020). Still, *wild* polyploid systems have lagged behind – even though they are essential for understanding the ecological context and evolutionary mechanisms of polyploid formation, establishment, and persistence (Soltis et al., 2016).

The allotetraploid monkeyflower, *Mimulus sookensis*, is one such system, having formed recently through hybridization between diploid species in the well-studied *M. guttatus* species complex (Benedict et al., 2012; Sweigart et al., 2008). Studies of allozyme (Benedict 1993) and nucleotide sequence variation (Modliszewski & Willis, 2012; Sweigart et al., 2008) have pointed to *M. guttatus* and *M. nasutus* as the diploid progenitors of *M. sookensis*. These two closely related diploid species range across much of western North America (Figure S1), and in areas of secondary sympatry, historical and contemporary introgression is common – largely from *M. nasutus* into *M. guttatus* (Brandvain et al., 2014; Kenney & Sweigart, 2016; Sweigart & Willis, 2003; Zuellig & Sweigart, 2018). The two diploid species also overlap with *M. sookensis*, which has been sampled from Vancouver Island, British Columbia, Canada to California, USA (Figure S1).

Although the geographic overlap and ongoing hybridization of *M. guttatus* and *M. nasutus* seem to build a strong case for their status as progenitors of *M. sookensis*, several key questions about the evolutionary origin of this allotetraploid species remain. One is the number of times it has evolved. Using nucleotide sequence polymorphism to investigate genetic variation in natural populations of *M. sookensis*, an early study with only two nuclear loci suggested the species had at least two evolutionary origins (Sweigart et al., 2008). A few years later, with an additional six nuclear loci and three chloroplast loci, the estimate increased to 11 independent origins (Modliszewski & Willis, 2012). Paradoxically, the same study showed no evidence of postzygotic reproductive isolation among any *M. sookensis* lines, which might be expected with multiple origins if duplicate gene copies are subject to divergent histories of degenerative mutation (Lynch & Force, 2000). Another question is the timing of the allotetraploid’s evolutionary origin. With only a handful of loci, previous studies have not attempted to estimate when *M. sookensis* evolved.

A final outstanding question is whether *M. guttatus* is a true progenitor. Although support for *M. nasutus* as a progenitor is strong, evidence for *M. guttatus* is mixed. Phenotypically, *M. sookensis* is nearly identical to *M. nasutus* (Benedict et al., 2012); both species are highly selfing, with small, often cleistogamous flowers. *M. guttatus*, on the other hand, is a large-flowered, primarily outcrossing species, and F1 hybrids from crosses with *M. nasutus* show dominance in floral traits toward *M. guttatus* (Fishman et al., 2002). Genetically, too, there is a much closer match between *M. nasutus* and *M. sookensis*, with molecular studies invariably showing high sequence similarity between one *M. sookensis* gene copy and *M. nasutus* alleles (Benedict 1993; Modliszewski & Willis, 2012; Sweigart et al., 2008). Although these same studies also show sequence similarity between the alternative *M. sookensis* gene copies and *M. guttatus* alleles, the exceptionally high genetic diversity of *M. guttatus* (Brandvain et al., 2014; Puzey et al., 2017) makes it nearly impossible to identify a direct progenitor. Even *M. nasutus* carries only a subset of the genetic variation present in *M. guttatus*, and the two species have very few fixed differences (Brandvain et al., 2014; Sweigart & Willis, 2003). In addition to *M. guttatus* and *M. nasutus*, the species complex includes several other closely related, often interfertile species (Vickery, 1978). Most of these species are unlikely progenitors, as they have restricted geographic distributions well outside the range of *M. sookensis*. However, one of them – *M. decorus* – is an outcrossing species with a range that largely overlaps both *M. nasutus* and *M. sookensis* (Figure S1).

In this study, we generate and leverage a new, chromosome-scale reference genome assembly for *M. sookensis* to clarify the details of its evolutionary history. Using whole genome sequence data (WGS) from *M. sookensis* accessions collected from across the species range, we perform population genomic analyses to determine the number and timing of evolutionary origins. We also revisit the question of ancestry, taking advantage of published WGS data from diverse *M. guttatus* complex accessions to identify the likely progenitors of *M. sookensis*. With these new genomic resources for *M. sookensis*, we expect this species to become a model for understanding the ecological context and genetic mechanisms of polyploid evolution.

## Methods

### Plant material, molecular methods, and sequencing

To generate plant material for the *M. sookensis* reference assembly, we grew a single plant from selfed seed of FAN36, an accession originally collected on Vancouver Island in British Columbia, Canada (Table S1). Because wild *M. sookensis* is primarily selfing, the FAN36 line is highly inbred and homozygous. This plant was grown in the University of Georgia Botany greenhouses in a 4-in pot with moist Fafard 4P growing mix (Sun Gro Horticulture, Agawam, MA) under 16-h days at 23°C and 8-h nights at 16°C. Once the plant had grown to a large size with several branches, we shipped it to Phase Genomics (Seattle, WA) for high-molecular weight DNA extraction, PacBio sequencing, and Hi-C chromatin capture.

Phase Genomics used ∼250 mg of leaf tissue from FAN36 to perform DNA extraction with the Qiagen MagAttract HMW DNA kit. Libraries were prepared using a PacBio Sequel II Binding Kit 2.2 and sequenced on a PacBio Sequel IIe instrument, producing 1,639,773 CCS HiFi reads with a total length of 24.27 Gb (average read length: 14.8 kb). Chromatin conformation capture data were generated using the Phase Genomics Proximo Hi-C 3.0 Kit, which is a commercially available version of the Hi-C protocol (Lieberman-Aiden et al., 2009). The resulting library was sequenced on an Illumina HiSeq 4000, generating a total of 242,673,575 150-bp PE reads.

We used new and previously generated whole genome sequence (WGS) data from 11 wild *M. sookensis* accessions collected from 10 populations across the species range (Table S1). Selfed seeds for each accession were grown as for FAN36 above. We performed DNA extractions using a CTAB-chloroform protocol (Fishman 2020) on bud and leaf tissue. DNA was submitted to Duke Center for Genomic and Computational Biology for sequencing where 150-bp PE reads were generated on an Illumina NovaSeq 6000 S1 lane.

### Genome assembly and annotation

To generate contigs for the *M. sookensis* assembly, we assembled the FAN36 HiFi reads with hifiasm v0.19.3 (Cheng et al., 2021) using the -l0 parameter to skip purging and preserve homoeologous regions. This method resulted in an initial assembly of 748 Mb with 3,671 contigs (N50 = 0.96 Mb). Evaluation of this assembly with KRAKEN v2.1.1(Wood et al., 2019) showed evidence of microbial contamination. Subsequent investigation with ProxiMeta (Press et al., n.d.; Stewart et al., 2018) to perform metagenome deconvolution, CheckM, v1.0.11 (Parks et al., 2015) to assess quality, and Mash v1.1.1 (Ondov et al., 2016) to determine taxon identity, revealed the inclusion of 32 microbial genome clusters. Removing all prokaryotic sequences from the initial contig assembly resulted in a draft assembly of 590 Mb with 979 contigs (N50 = 1.19 Mb).

To generate chromosome-scale scaffolds, we mapped Hi-C reads to the draft assembly following the Phase Genomics Proximo Hi-C Kit recommendations (https://phasegenomics.github.io/2019/09/19/hic-alignment-and-qc.html). Briefly, we aligned reads using BWA-MEM v0.7.17 (Li & Durbin, 2010) with the -5SP and -t 8 options specified and all other options default. We flagged PCR duplicates (later excluded from the analysis) using SAMBLASTER v0.7.17 (Faust & Hall, 2014) and filtered alignments using SAMtools v1.9 (Danecek et al., 2021) with the -F 2304 filtering flag to remove non-primary and secondary alignments. To create scaffolds, we used a single-phase procedure first used in Bickhart et al., 2017. As in the LACHESIS method (Burton et al., 2013), this procedure computes a contact frequency matrix from the aligned Hi-C read pairs (normalized by the number of Sau3AI restriction sites on each contig) and constructs scaffolds to optimize expected contact frequency in Hi-C data. We used Proximo to perform ∼20,000 iterations of this dataset to optimize the number of scaffolds, as well as the orientation and order of contigs (maximizing concordance with the observed Hi-C data). Using Juicebox v0.7.17 (Durand et al., 2016; Rao et al., 2014) to manually correct scaffolding errors produced a final, 574-Mb *M. sookensis* FAN v1 assembly that includes 28 chromosome-scale scaffolds ranging in size from 10.18 Mb to 26.69 Mb (the assembly also includes a 5-Mb scaffold with sequences that could not be assembled into the main chromosome-level scaffolds.). A 15 Mb fungal contaminant identified via NCBI BLAST (blast.ncbi.nlm.nih.gov, Altschup et al., 1990) was removed from the assembly. We used BUSCO v5.2.2 (Simão et al., 2015) in genome mode with eukaryota odb10 to evaluate the completeness of the assembly.

To generate a preliminary annotation for the *M. sookensis* FAN v1 reference, we used Liftoff v1.6.3 (Shumate & Salzberg, 2021) and the *M. guttatus* v3 annotation (phytozome.org). Because *M. sookensis* should contain twice the number of genes as its diploid progenitor, we used the -*copies* flag along with *-sc 0.75* to ensure high quality lift over.

### Subgenome identification

To identify homeologous chromosome pairs, we performed genome comparisons between *M. sookensis* and its presumed diploid progenitors. We used MiniMap2 v2.26 (Li, 2018) to compare the *M. sookensis* FAN v1 genome to the *M. guttatus* IM62 v3 and *M. nasutus* SF v1 reference assemblies (phytozome.org) and generated dot plots using dotPlotly (Tom Poorton, https://github.com/tpoorten/dotPlotly).

We assigned subgenomes by mapping previously generated WGS from *M. nasutus* SF (Table S1) to the *M. sookensis* FAN v1 assembly. We used TRIMMOMATIC v0.36 (Bolger et al., 2014) to remove adapters, BWA-mem v0.7.17 (Li, 2013; Li & Durbin, 2010) to align SF to FAN v1, and Picard v2.27.4 (http://broadinstitute.github.io/picard) to add read groups and remove optical and PCR duplicates. We then filtered reads for MQ > 28 in SAMtools v1.16.1 (Danecek et al., 2021). To calculate SF *M. nasutus* read coverage across the *M. sookensis* FAN v1 assembly, we used the coverage tool in SAMtools (Danecek et al., 2021). Additionally, to assess SF-FAN sequence similarity, we generated a VCF file using the mpileup and call tools from BCFtools v1.15.1(Danecek et al., 2021). We used VCFtools v0.1.16 (Danecek et al., 2011) to remove indels and include only biallelic sites with quality > 29. We filtered each VCF to include only sites with read depth ≥10 and <mean+2*stdev. We used Pixy v1.27.beta1 (Korunes & Samuk, 2021) to calculate nucleotide diversity (*π*) between FAN and SF.

We used MiniMap2 v2.26 and Circos v0.69 (Krzywinski et al., 2009) to visualize regions of homeology in the *M. sookensis* genome. To explore synteny across the genome between *M. sookensis* and putative progenitors, we input peptide fasta and gff files for *M. sookensis* FAN v1, *M. guttatus* IM62 v3, and *M. nasutus* SF v1 into GENESPACE v1.2.3 (Lovell et al., 2022).

### Population genomic analyses

To investigate patterns of natural variation in *M. sookensis*, we aligned WGS data from 11 wild accessions (Table S1) to the *M. sookensis* FAN v1 reference assembly using the methods described above for SF *M. nasutus*. Using the mpileup and call tools from BCFtools v1.15.1 (Danecek et al., 2021), we generated a VCF for each accession. We used VCFtools v0.1.16 (Danecek et al., 2011) to remove indels and include only biallelic sites with quality > 29. We filtered each VCF to include only sites with read depth ≥10 and <mean+2*stdev and 80% of individuals were genotyped. We performed a principal component analysis (PCA) in R [4.1.3] (R core team 2021) using the package “SNPRelate” (Zheng et al., 2012). To visualize the axes, we used ggplot2 (Wickham H, 2016) in R. We used Pixy v1.27.beta1 (Korunes & Samuk, 2021) to calculate heterozygosity and *π*. For these and all downstream analyses, we took advantage of having Illumina WGS data from the same FAN36 accession used to generate the reference assembly. To minimize problematic alignments caused by assembly errors, we excluded reads from any of the 11 wild *M. sookensis* lines that mapped to genomic regions where FAN36 itself was identified as heterozygous.

To perform genome comparisons between *M. sookensis* and its potential diploid progenitors, we separated *M. sookensis* into its two subgenomes. We aligned the 11 wild accessions to the FAN v1 reference using BWA-mem v0.7.17 (Li & Durbin, 2010), split resulting BAM files by chromosome, and merged the split BAM files using SAMtools v1.16.1 merge (Danecek et al., 2021) into separate files representative of the subgenomes. We then converted BAM files back to the FASTQ format using SAMtools v1.16.1 bam2fq (Danecek et al., 2021). We aligned the separate subgenomes of 11 *M. sookensis* to the *M. guttatus* v5 reference assembly, along with previously generated WGS from *M. guttatus*, *M. nasutus*, *M. decorus*, and a *M. dentilobus* accession as an outgroup (Table S1). Using GATK v3.8-1 HaplotypeCaller with the -allSites flag (McKenna et al., 2010) we generated a VCF. We removed indels and used only biallelic sites with read depth ≥10 and < mean+2*stdev that passed the following filters: mapping quality (MQ) > 30, mapping quality rank sum (MQRankSum) > - 12.5, fisher strand (FS) < 60, quality depth (QD) > 2, quality (QUAL) > 30 and read position rank sum (ReadPosRankSum) > -8. Additionally, we filtered individual VCF files for each sample to include only sites with read depth ≥ 3 and < mean+2*stdev. We used VCFtools v0.1.16 (Danecek et al., 2011) to remove sites with > 20% missing data.

From this dataset, we generated neighbor joining (NJ) and maximum likelihood (ML) trees that included all 22 *M. sookensis* subgenomes, as well as closely related diploid species in the *M. guttatus* complex to investigate the evolutionary history of *M. sookensis*. In R, we used the package “ape” to construct the NJ tree and “phangorn” to perform 1000 bootstraps on a downsampled dataset of 14,000 SNPs. After conversion into PHYLIP format with custom scripts from Simon Martin (https://github.com/simonhmartin/genomics_general/tree/master), we generated the ML tree with the full dataset using IQ-Tree v2.2.0 (Nguyen et al., 2015) with TVM+F+R2 as the best fit model. The ML tree included 3,576,989 (1,711,738 parsimony informative) sites. Finally, we identified fourfold degenerate sites using a custom script (Tim Sackton: https://github.com/tsackton/linked-selection/tree/master/misc_scripts) and then used a python script (Note S1 in Garner et al., 2016) to calculate nucleotide diversity (*π_sil_*) and divergence (*d_xy_*) among samples.

To characterize variation in species relationships across the genome, we used IQ-Tree v2.2.0 to generate 16,754 gene trees including only genes with variant calls for all taxa. For each gene, we used BCFtools v1.15.1(Danecek et al., 2021) consensus to generate a concatenated fasta and the seqtk v1.3 ‘randbase’ tool (https://github.com/lh3/seqtk) to randomly select one allele at heterozygous sites. Next, we used TWISST v0.2 (Martin & Van Belleghem, 2017) in two separate analyses to quantify support for particular (rooted) gene tree topologies given a specified set of species in the dataset. In the first analysis, we excluded *M. sookensis* samples and grouped the remaining diploids into three “species”: *M. decorus*, northern *M. guttatus*, and southern *M. guttatus* + *M. nasutus*. We quantified support for each of the three possible gene tree topologies with *M. dentilobus* as the outgroup. Our rationale for combining southern *M. guttatus* and *M. nasutus* into one “species” and for separating *M. guttatus* into two was that these groupings form well-supported clades with whole genome data (Brandvain et al., 2014, also see Results). Moreover, because our intent with this analysis was to investigate whether discordant gene trees might help explain conflicting whole-genome tree topologies that have alternatively placed *M. decorus* within northern *M. guttatus* (Coughlan et al., 2020; Ivey et al., 2023) or sister to all of *M. guttatus* (Coughlan and Willis 2019), it was necessary to distinguish between geographic clades. In the second TWISST analysis, we included *M. sookensis* subgenomes and quantified support for one or more diploid relatives as a likely progenitor. In this analysis, we categorized the 105 rooted gene tree topologies (for five taxa plus *M. dentilobus* as an outgroup) into simplified “classes” defined by sister groupings between a focal *M. sookensis* subgenome and the diploid taxa. We also visualized genome-wide support for the most frequent of these topology classes by averaging topology weights in non-overlapping nine-gene windows.

To attempt to identify the maternal progenitor of *M. sookensis*, we characterized patterns of variation in the chloroplast and mitochondrial genomes. Using BWA-mem v07.17 (Li & Durbin, 2010) we aligned WGS from 11 *M. sookensis, 20 M. guttatus, 8 M. decorus, and 5 M. nasutus* to an *M. guttatus* IM62 mitochondria assembly (Mower et al., 2012), excluding regions annotated as derived from the chloroplast. We also aligned the same WGS data to the chloroplast genome of the *M. guttatus* IM767 v1 reference assembly (phytozome.org). Using the mpileup and call tools from BCFtools v1.15.1(Danecek et al., 2021), we generated a combined VCF for each genome and used VCFtools v0.1.16 (Danecek et al., 2011) to remove indels and include only biallelic sites with quality > 29. We excluded sites with heterozygous calls or with any missing data. For each organellar genome, we constructed a haplotype network using PopArt (Leigh & Bryant, 2015). The mitochondrial network used 233 sites (100 parsimony informative) and the chloroplast network used 181 sites (91 parsimony informative).

## Results

### Mimulus sookensis genome assembly

Consistent with its status as an allotetraploid, the *M. sookensis* genome assembly is 574 Mb, roughly twice the size of the reference genomes of closely related diploids (*M. guttatus* IM62 v3 = 339.2 Mb, *M. nasutus* SF v3 = 312.8 Mb, phytozome.org). Additionally, we recovered 28 chromosome-scale scaffolds in *M. sookensis* (Figure S2), double the base number of chromosomes in the group (Mukherjee & Vickery, 1962). Both synteny mapping and homology between *M. sookensis* and its diploid relatives show that each of the 14 chromosomes in *M. nasutus* and *M. guttatus* corresponds to exactly two chromosomes in *M. sookensis* (Figure 1, Figure S3). This sequence homology spans chromosomes end-to-end indicating there have been no large-scale deletions or chromosome loss in *M. sookensis*. We do find evidence of a *M. sookensis* specific inversion at the tip of chromosome 14 of subgenome A. Finally, BUSCO analysis revealed the new *M. sookensis* FAN v1 assembly to be 98.2% complete and, consistent with a recent whole genome duplication, 90.2% of BUSCOs are duplicated (C:98.4% [S:8.2%, D:90.2%], F:0.5%, M:1.1%, n:2326).

**Figure 1:**
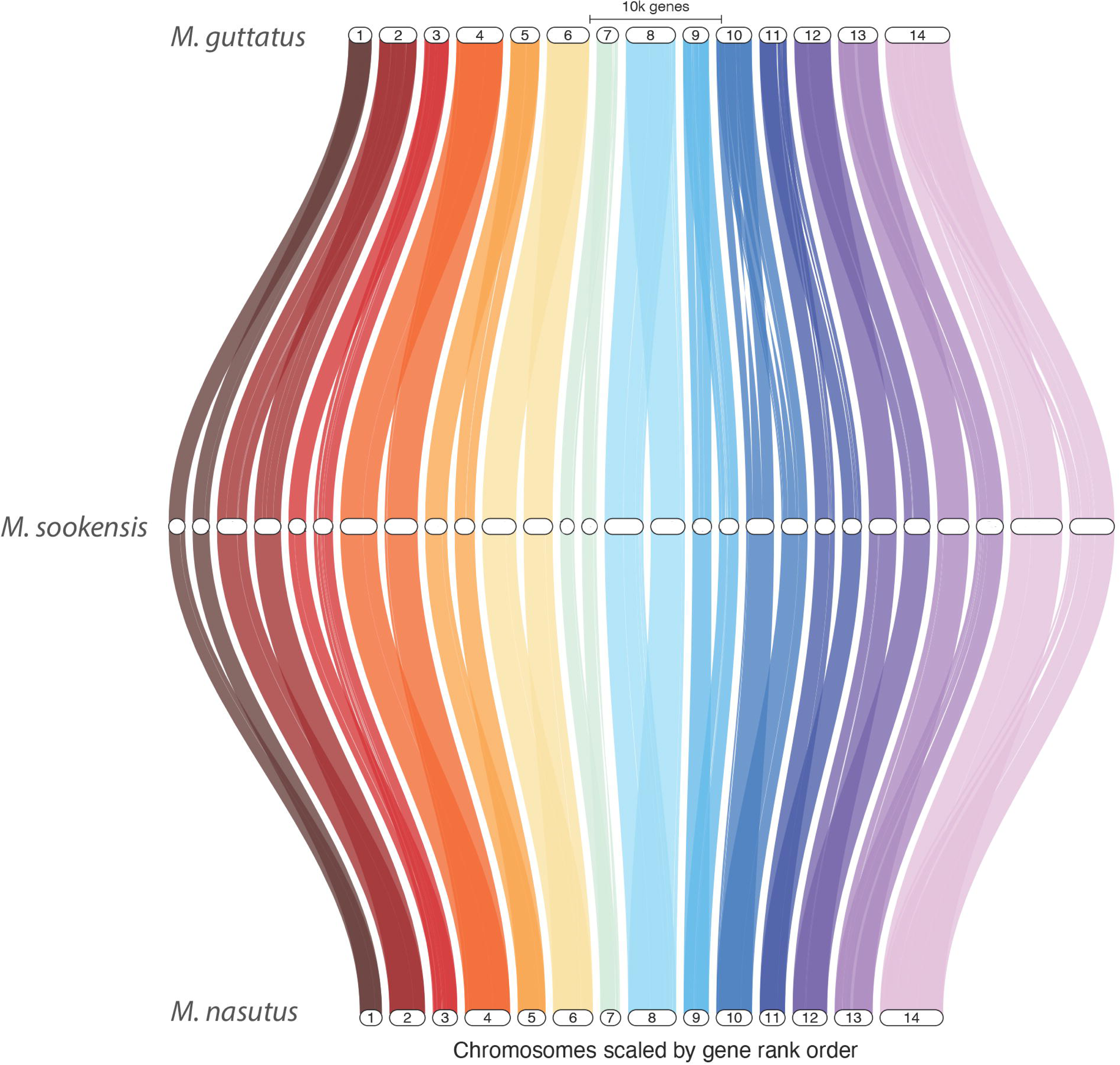
*M. sookensis* shows high levels of synteny with diploid relatives *M. nasutus* and *M. guttatus.* Genespace syntenic map created using the *M. sookensis* v1, *M. guttatus* v3, and *M. nasutus* v1 reference genomes.

### Subgenome identification

To define subgenomes, we aligned WGS from *M. nasutus* SF to the *M. sookensis* FAN v1 reference, reasoning that sequence from this putative progenitor should match only one set of 14 chromosomes. Indeed, for each pair of *M. sookensis* homeologs, *M. nasutus* read depth was much higher on one chromosome than on the other (Table 1). Using this analysis, we defined the set of 14 chromosomes with higher *M. nasutus* coverage as subgenome A (average read depth = 6.2) and the set with lower *M. nasutus* coverage as subgenome B (average read depth = 0.55). Nucleotide diversity is much lower when *M. nasutus* is compared to subgenome A (*π* = 0.005) than subgenome B (*π* = 0.025) and is also similar to nucleotide diversity within *M. nasutus* (*π* = 0.014: Garner et al., 2016). Taken together, these results point to *M. nasutus* as the progenitor of subgenome A but not subgenome B. The two subgenomes show little to no sequence synteny outside homeologous pairs (Figure S4).

### Evolutionary origin of sookensis

Despite sampling *M. sookensis* from locations spanning most of its range, we detected little population structure in this species with PCA revealing no distinction between northern and southern populations (Figure 2). Additionally, we discovered very little genetic variation in *M. sookensis*: average pairwise nucleotide diversity (*π*) is only 0.0045 ∼half that of its selfing progenitor (*M. nasutus*). These results are strong evidence of a single evolutionary origin of *M. sookensis*.

**Figure 2:**
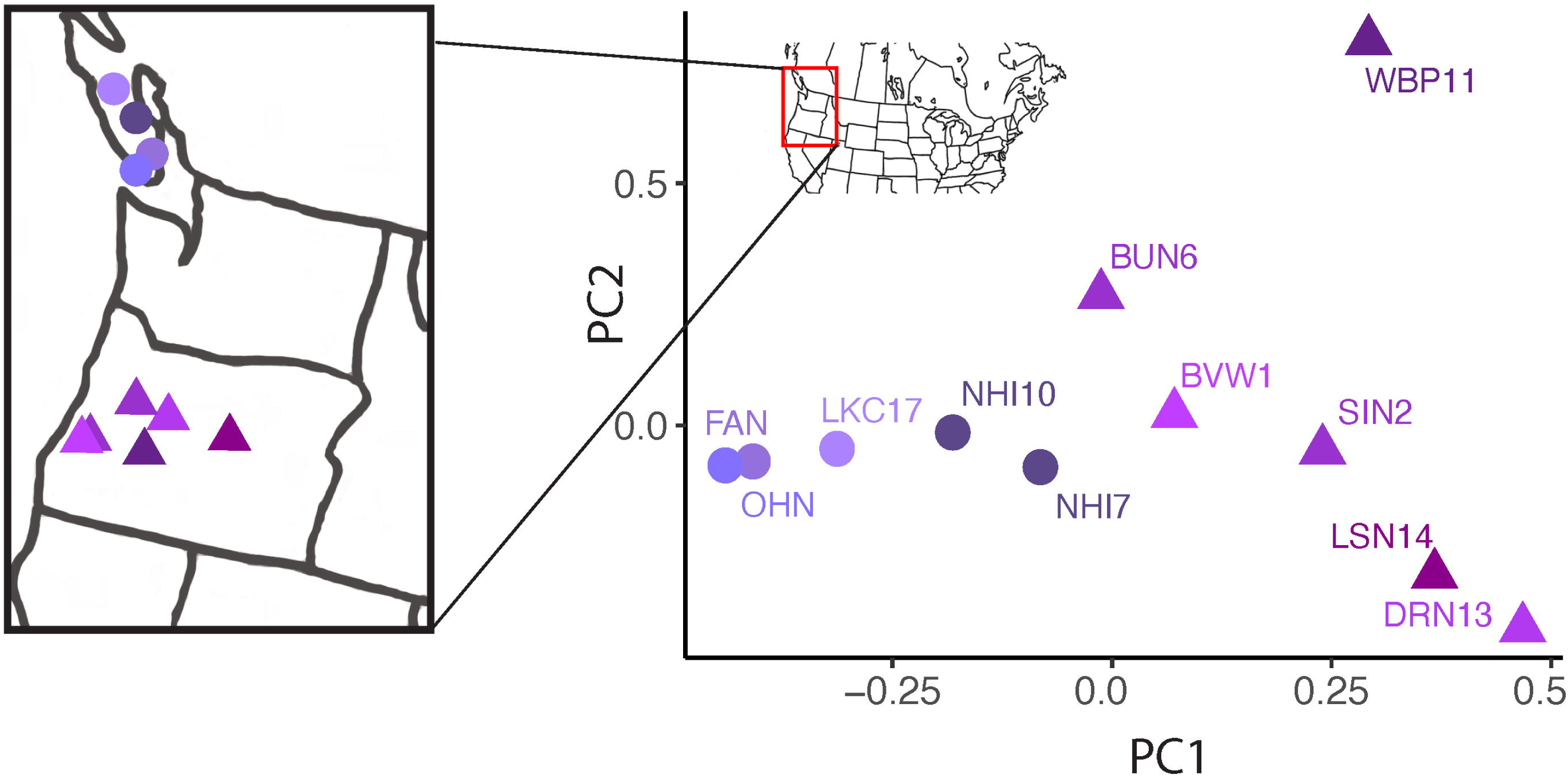
*M. sookensis* populations show little genetic structure in a PCA. PC1 and PC2 explain 19.21% and 15.01% of the variation respectively. Samples collected from Vancouver Island, BC, Canada are shown by circles and those collected from Oregon, USA are shown by triangles. Sample locations are shown in map on left (note one population has two accessions).

To investigate this origin more deeply, we separated *M. sookensis* into its two subgenomes for comparisons with diploid relatives in the *M. guttatus* complex. Consistent with a single allopolyploid origin, the two *M. sookensis* subgenomes form distinct monophyletic groups in maximum likelihood (Figure 3, Figure S6) and neighbor-joining (Figure S5) trees. The subgenome A samples are nested within the *M. nasutus* clade, confirming this species as a progenitor of *M. sookensis*. The identity of the other progenitor, however, is less certain. In contrast to previous studies (Modliszewski & Willis, 2012; Sweigart et al., 2008), which focused exclusively on *M. nasutus* and *M. guttatus* as potential progenitors (and did not include analyses of any other diploid relatives), we discovered that subgenome B clusters most closely with *M. decorus* (Figure 3, Figure S5). However, unlike with subgenome A, we did not observe the nested pattern of variation expected for a direct progenitor. We note there is extensive hybridization and incomplete lineage sorting among closely related members of the *M. guttatus* complex (Brandvain et al., 2014), which complicates phylogenetic reconstructions, and *M. decorus* has alternatively been placed within the northern *M. guttatus* clade (Coughlan et al., 2021; Ivey et al., 2023) and outside of *M. guttatus* altogether (this study, Coughlan & Willis, 2019). Even with this uncertainty about the identity of one progenitor, the pattern of variation – with each subgenome forming a well-supported, distinct clade – provides unambiguous support for a single evolutionary origin of *M. sookensis*. Moreover, assuming this origin arose through a single F1 individual, we can estimate it occurred only ∼71,000 years ago (with time of origin *T* ∼ *π*_sil_/*μ* and *π*_sil_ = 0.011 = average *π*_sil_ of subgenome A and B [Table S2], assuming *μ* = 1.5×10^-8^ following Brandvain et al., 2014).

**Figure 3:**
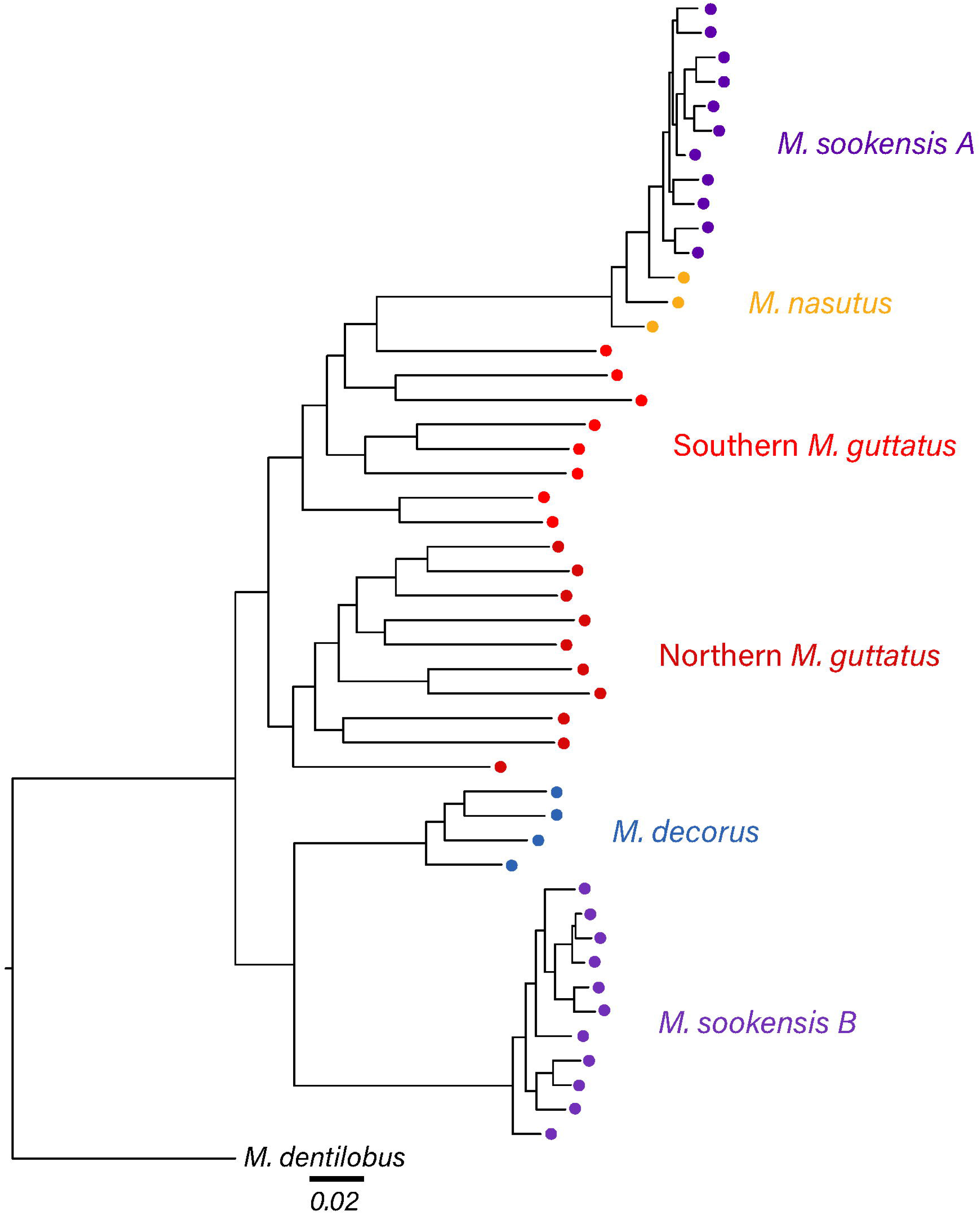
Maximum likelihood tree of *M. sookensis* subgenomes and diploid relatives in the *M. guttatus* species complex. The ML tree was generated in IQ Tree using the TVM + F + R2 model from 1,711,738 parsimony informative sites with *M. dentilobus* as an outgroup.

We next used TWISST (Martin & Van Belleghem, 2017) to quantify support for alternative evolutionary histories across the genome. In an initial analysis excluding *M. sookensis* (Figure S7), we discovered that gene tree topologies with *M. decorus* outside of *M. guttatus* are only slightly more common (∼43%) than topologies with *M. decorus* sister to northern *M. guttatus* (∼33%). Next, in an analysis with the full taxon dataset, we quantified support for one or more diploid species as the closest relative of *M. sookensis* subgenome B. We defined five “species” in this analysis (in addition to *M. dentilobus* as the outgroup): *M. sookensis* subgenome B, *M. decorus*, northern *M. guttatus*, southern *M. guttatus*, and *M. nasutus* + *M. sookensis* subgenome A. Our rationale for collapsing *M. nasutus* and subgenome A into a single species was that these samples form a well-supported clade in the whole-genome tree (Figure 3) and it reduces the number of tree topologies to a manageable number. We categorized the 105 rooted gene tree topologies into simplified classes defined by sister groupings between *M. sookensis* subgenome B and all possible combinations of the four other diploid species (Figure 4, Figure S8). The most common class of topologies have *M. decorus* as sister to subgenome B (∼36%), mirroring the whole-genome species tree (Figure 3). This relationship occurs more than twice as often as the second most common class of topologies (subgenome B equally related to *M. decorus*, *M. guttatus*, and *M. nasutus*: ∼15%) and more than three times the third most common class (subgenome B sister to northern *M. guttatus*: ∼11%). Across the genome, there is alternating support for these top three topology classes (Figure S9), with little evidence of recent introgression (i.e., large contiguous blocks supporting a particular topology) from any of these diploid species into *M. sookensis*. The fourth most common topology class (∼8%) places subgenome B as sister to the *M. nasutus*/subgenome A group (Figure 4, Figure S10), which might be due to gene conversion and/or homeologous recombination between the *M. sookensis* subgenomes.

**Figure 4:**
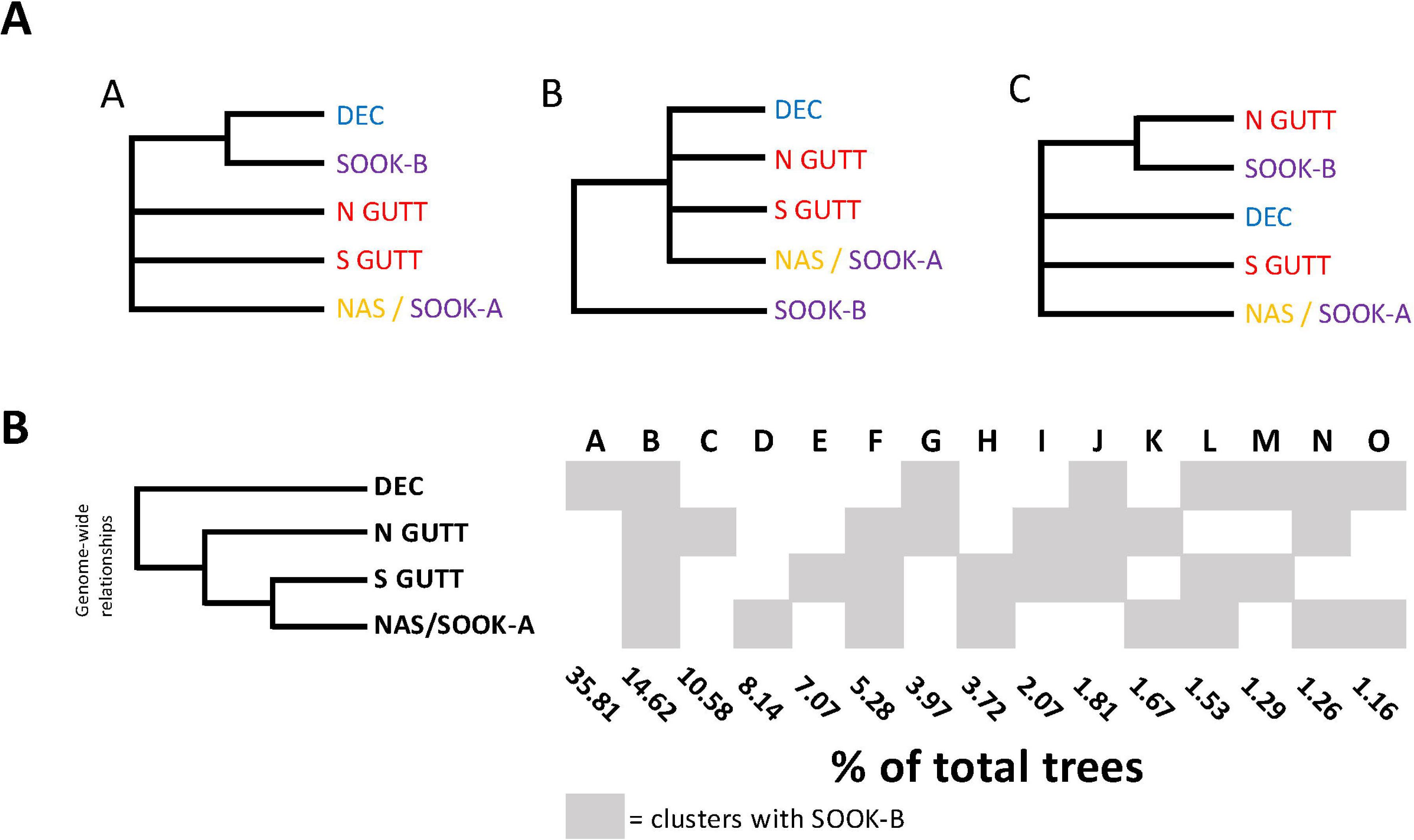
Genome-wide TWISST weightings quantifying support for one or more diploid species as the closest relative to *M. sookensis* subgenome B. Data are from 16,754 rooted gene trees with five “species” (subgenome B, *M. decorus*, northern *M. guttatus*, southern *M. guttatus,* and subgenome A + *M. nasutus*) plus *M. dentilobus* (not shown) as the outgroup. (A) The 105 topologies were grouped into 15 simplified classes (A-O) defined by which diploid species (or combination) is most closely related (i.e., sister) to *M. sookensis* subgenome B (indicated with gray shading). Frequencies of each topology class are given to the right of the table; the whole-genome species tree is depicted at the top. (B) The top three topology classes.

Finally, in an attempt to identify the maternal progenitor of *M. sookensis*, we analyzed patterns of variation in mitochondrial and chloroplast genomes. Although previous analyses with fewer loci and taxa found no species-diagnostic haplotypes in the *M. guttatus* complex (Modliszewski & Willis, 2012; Vallejo-Marín et al., 2016), both organellar genomes in this study revealed structure between species. In both the mtDNA and cpDNA haplotype networks, *M. nasutus* and *M. sookensis* cluster together in a group that also includes four *M. guttatus* samples (Figure 5). Taken at face value, this result might suggest that shared variation between diploid relatives precludes identification of the *M. sookensis* maternal progenitor. However, three of the four *M. guttatus* accessions in this group are from populations with ongoing introgression from *M. nasutus* (Brandvain et al., 2014; Kenney & Sweigart, 2016). This fact, together with the finding that almost all *M. decorus* samples cluster separately (except for one sample – LL – that groups with *M. sookensis* only in the mtDNA network) suggests that *M. nasutus* is the maternal progenitor of *M. sookensis*.

**Figure 5:**
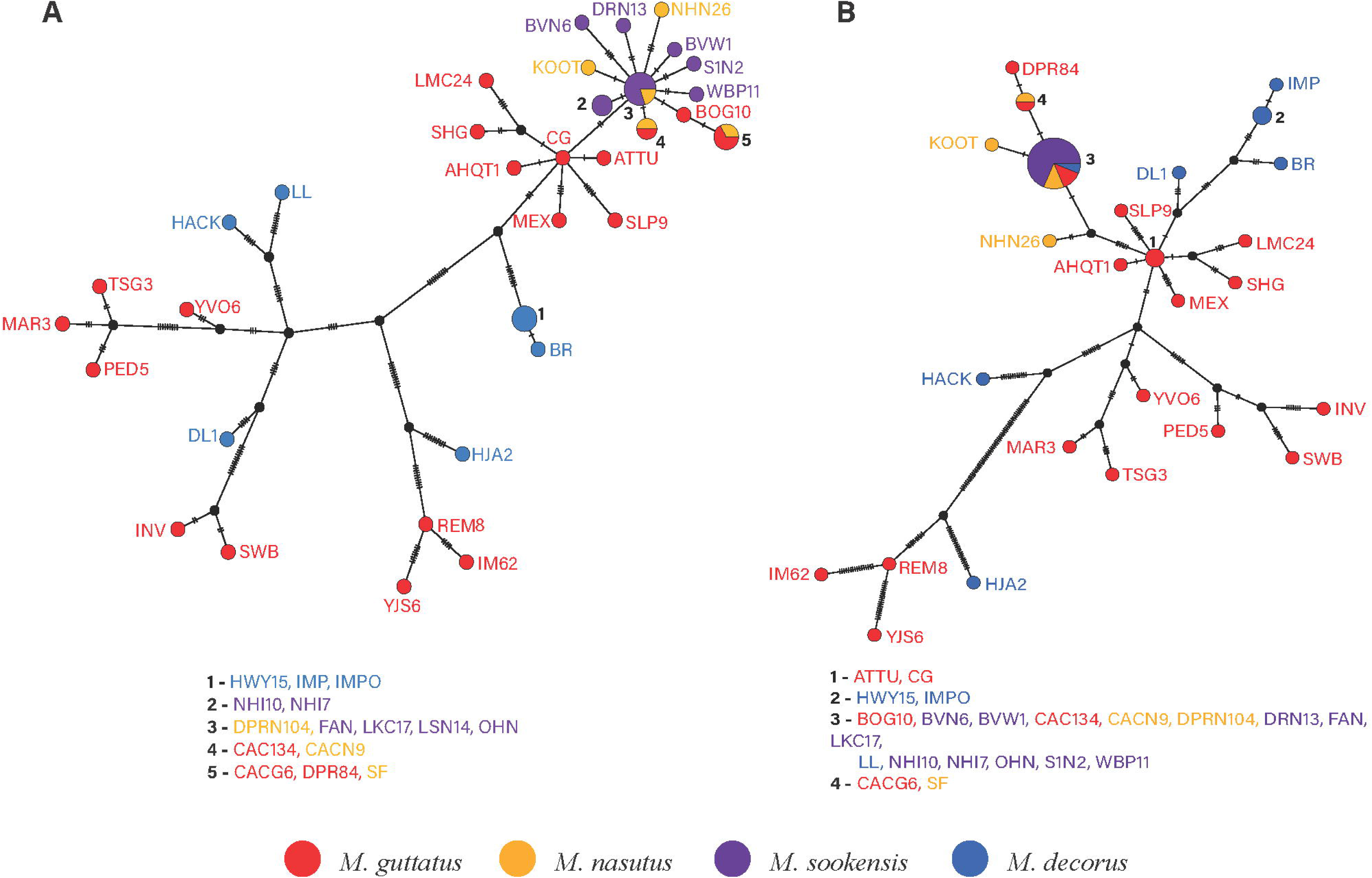
Haplotype networks of organellar genomic variation point to *M. nasutus* as the maternal progenitor of *M. sookensis*. (A) Chloroplast haplotype network generated using 91 parsimony-informative sites. (B) Mitochondrial haplotype network generated using 100 parsimony-informative sites. Haplotypes represented by more than one sample are indicated with numbers (individual samples in numbered groups are listed below each network).

## Discussion

Understanding why polyploidy is so ubiquitous across the angiosperm phylogeny requires a focus on wild systems. Here, we generate a chromosome-scale reference genome assembly for the wild allotetraploid *Mimulus sookensis*, which formed through hybridization between closely related yellow monkeyflower species in the *M. guttatus* complex. This new genome assembly allowed us to interrogate the evolutionary history of *M. sookensis*, providing definitive evidence of a single origin of the species, which we estimate occurred ∼71,000 years ago. Additionally, our analysis defined the two distinct subgenomes of *M. sookensis*, one of which is clearly derived from the diploid selfing species, *M. nasutus*. Going forward, this new genome will be an invaluable resource for understanding the genetic mechanisms of adaptation in *M. sookensis*, as well as divergence from its progenitors.

### Single origin of M. sookensis

Our discovery that *M. sookensis* has a single evolutionary origin differs dramatically from previous work suggesting as many as 11 origins (Modliszewski & Willis, 2012). How do we explain this large discrepancy? Inference in the earlier study was based on nucleotide sequence variation at a much more limited number of loci (8 nuclear, 3 chloroplast), only two of which showed truly high levels of polymorphism consistent with multiple origins (Modliszewski & Willis, 2012). In light of our genome-wide data, which clearly show low variability in *M. sookensis* across most of the genome (*π* = 0.0045), these two loci appear to be outliers. Indeed, we reexamined sequence variation at these two loci (*Mg1* and *Mg5*: Modliszewski & Willis, 2012) in our WGS dataset and found the same unusually high levels of polymorphism. This cause of high variation at these two loci is unknown but could be due to hybridization with other polyploid relatives in the *M. guttatus* complex (see below).

Like in *M. sookensis*, the availability of genome-wide sequence data and/or reference genomes has recently overturned or clarified the origin stories of several other wild and domesticated polyploid species. A high-quality reference genome for *Capsella bursa-pastoris*, for instance, helped uncover evidence of a homeologous exchange event that appears to have evolved very early in the species’ evolutionary history (the exchange is geographically widespread and present in all studied accessions), suggesting that one origin with subsequent introgression from diploid progenitors might be more likely than multiple origins (Penin et al., 2023). In *Arabidopsis suecica*, the story is reversed with multiple origins now inferred from extensive shared variation with its diploid progenitors (Novikova et al., 2018), instead of the single origin suggested by earlier studies of microsatellite variation (Jakobsson et al., 2006) In crops too, recent genomic analysis has begun to clarify what are often very complex evolutionary origins involving multiple rounds of hybridization between several different progenitors (e.g., bread wheat: Liu et al., 2022; Pont et al., 2019).

For several reasons, the finding of a single evolutionary origin of *M. sookensis* is somewhat surprising. One is simply a lack of precedent: among the wild polyploid species that have been studied, multiple origins seem to be the rule (Soltis et al., 2014, 2016; Soltis & Soltis, 1999; Another is that there might be ample opportunity for polyploid formation, with considerable historical and contemporary hybridization between some diploid species of the *M. guttatus* complex (Brandvain et al., 2014; Sweigart & Willis, 2003). Given the recent origin of *M. sookensis* and its fairly wide geographic distribution (at least from northern California to southern British Columbia), the hypothesis of multiple origins seems entirely plausible. On the other hand, that fact that *M. sookensis* was immediately able to self-fertilize when formed (one progenitor – *M. nasutus* – is a selfer and all *Mimulus* species are self-compatible) was likely a huge boon to its establishment (Fowler & Levin, 2016; Soltis et al., 2014) and might also have aided in its spread (Razanajatovo et al., 2016; Wright et al., 2013).

### Search for the diploid progenitors

In line with previous evidence (Benedict et al., 2012; Sweigart et al., 2008), our study clearly identifies *M. nasutus* as a progenitor of *M. sookensis*. With the new *M. sookensis* FAN genome assembly in hand, we were able to identify the set of 14 chromosomes derived from *M. nasutus*, which we call subgenome A. Additionally, by analyzing variants across the entire mitochondrial and chloroplast genomes, our study provides the strongest evidence to date of *M. nasutus* as the maternal progenitor. Due to extensive shared variation in the *M. guttatus* complex, previous studies of these genomes, which included either fewer loci (Modliszewski & Willis, 2012) or taxa (Vallejo-Marín et al., 2016), provided little resolution to distinguish among potential maternal progenitors. In this study, however, we find a consistent group of taxa that clusters together in both organellar networks; the group includes all *M. sookensis* samples, all *M. nasutus* samples, and four *M. guttatus* samples (plus one *M. decorus* sample in the mitochondrial network). Although, at first glance, this pattern seems to suggest shared variation in the diploid progenitors, three of the four *M. guttatus* samples that cluster in this group are from populations sympatric with *M. nasutus*, where extensive directional introgression (from *M. nasutus* into *M. guttatus*) has been documented (Brandvain et al., 2014; Kenney & Sweigart, 2016; Sweigart & Willis, 2003). Thus, it appears that hybridization in these sympatric populations may have resulted in organellar capture as has recently been found in other *Mimulus* species (Nelson et al., 2021). Moreover, these results build on previous intuition that, given the strong triploid block between *M. sookensis* and its diploid relatives (Sweigart et al., 2008), initial formation likely occurred via unreduced gametes in an F1 hybrid with *M. nasutus* as the seed parent.

Despite our extensive population genomic sampling, the identity of the second progenitor to *M. sookensis* remains somewhat mysterious. Although subgenome B clusters most closely with *M. decorus*, we do not observe the nested pattern of variation expected for a direct progenitor. Casting further doubt on *M. decorus* ancestry, the new genome assembly indicates that neither of the *M. sookensis* homeologs to chromosome 8 has the *DIV1* inversion (Figure S3), which is carried by all perennial ecotypes of *M. guttatus* (Lowry & Willis, 2010; Oneal et al., 2014) and is also found in perennial *M. decorus* (Coughlan and Willis 2019). Patterns of sequence variation in the *DIV1* inversion suggest it is not new (Twyford and Friedman 2015) and its presence in *M. decorus* suggest it evolved prior to this species’ split with *M. guttatus* ∼230 KYA (Coughlan & Willis, 2019). Thus, the recent origin of *M. sookensis* ∼71 KYA seems to rule out as a progenitor any species fixed for the *DIV1* inversion.

It should be noted that *M. guttatus* remains a candidate progenitor, as this species is exceptionally diverse (Brandvain et al., 2014; Puzey et al., 2017) and has a large geographic range that overlaps with *M. sookensis* and *M. nasutus*. Moreover, in at least three populations, the three species currently co-occur (ROG and NHI; Sweigart et al., 2008; Catherine Creek, unpublished results). Still, the fact that subgenome B of *M. sookensis* clusters outside of the *M. guttatus* clade (Figure 3) suggests the polyploid carries ancestry either from unsampled *M. guttatus* lineages or from other close relatives in the species complex. This ancestry might have been present in *M. sookensis* from the start (derived from the original progenitor) or it might have been introduced later through hybridization with close relatives. High sequence similarity with *M. nasutus* across the entirety of subgenome A seems to suggest a primary influence of the former, as evidence of hybridization should be found in both subgenomes. Additionally, patterns of shared variation between subgenome B and its diploid relatives (*M. guttatus* and *M. decorus*) are relatively constant across the genome (Figure S9) instead of blocky as might be expected under a scenario of recent hybridization (Brandvain et al., 2014).

One intriguing possibility is that the unidentified progenitor could be another tetraploid in the *M. guttatus* complex. Although there are no other widely distributed polyploids in the complex, there have been reports of locally distributed *M. guttatus* autopolyploids (Vickery, 1995) and *M. decorus*-like polyploids (Coughlan et al. 2020), both of which overlap geographically with *M. sookensis* and *M. nasutus*. To investigate sequence similarity between *M. sookensis* and the *M. decorus*-like polyploid (which is a putative allotetraploid between *M. decorus* and an unknown species: J. Coughlan, pers. comm.), we aligned publicly available WGS from two accessions of this taxon (Table S1) to the *M. sookensis* FAN genome assembly. Contrary to the expectation if this *M. decorus*-like polyploid was a direct progenitor of subgenome B, we found similar sequence coverage across both *M. sookensis* subgenomes (A: 43.88%, B:56.69%). Future sampling efforts should target other, locally distributed polypoid taxa in the *M. guttatus* complex toward the goal of gaining additional insight into *M. sookensis* ancestry.

### Conclusions and future directions

Our analyses of population genomic variation in *M. sookensis* provide definitive evidence of the species’ recent and unique evolutionary origin. Although the direct progenitor of subgenome B remains somewhat mysterious, it is likely to have been an outcrosser, as other selfing species in the complex do not overlap with *M. sookensis*. One of the most salient features of *M. sookensis* is its nearly identical phenotype to *M. nasutus* (Benedict et al., 2012; Sweigart et al., 2008) and a key question in this system is what genetic mechanisms drive this similarity. Our finding of two distinct subgenomes corroborates previous studies of marker variation (Modliszewski & Willis, 2012; Sweigart et al., 2008), which have shown *M. sookensis* acts like a “fixed heterozygote,” carrying gene copies from each of its progenitors. Although these intact subgenomes suggest there has not been large-scale homoeologous recombination, we do find a signal of local genomic similarity between subgenomes A and B, with the two subgenomes clustering together in ∼8% of gene trees (Figure S10). This sequence similarity is distributed across much of the genome and might indicate a history of gene conversion, but whether this mechanism results in the fixation of *M. nasutus* alleles in *M. sookensis* or explains the phenotypic resemblance of these two species remains to be seen. The new *M. sookensis* genome assembly also reveals a putative inversion at the end of chromosome 14A (Fig S3), which might be unique to the allotetraploid given that it has not been seen in the genomes of its diploid relatives (phytozome.org). With the origin of *M. sookensis* now clarified, and a new genome assembly available for this wild polyploid model system, we are now poised to address longstanding questions about the genetic and evolutionary mechanisms of polyploid establishment and spread.

## Author Contributions

MRW and ALS designed and conceived the research. HM generated the assembly and Hi-C scaffolding and wrote the text related to these methods. MRW collected and analyzed the data. MRW and ALS wrote the manuscript.

## Data Accessibility

Whole genome sequence data of lines used will be deposited to the NCBI Sequence Read Archive upon acceptance. The *Mimulus sookensis* FAN v1 reference genome will be made publicly available.

## Acknowledgements

We thank J. Modliszewski and J. Willis who kindly provided seeds and shared whole genome sequence data for *M. sookensis*. We also thank J. Coughlan for sharing sequence data for *M. decorus*. We are grateful to J. Burke, J. Coughlan, K. Dyer, M. Farnitano, N. Gonzalez, J. Leebens-Mack, J. Modliszewski, S. Mantel, R. Schmitz, G. Sandstedt, VA. Sotola for helpful discussions. S. Mantel, M. Farnitano, and N. Gonzalez provided valuable feedback on the manuscript. This work was supported by an SSE Lewontin Award to MRW, a UGA Plant Center award to MRW, a Robin Hightower Genetics Graduate Support Fund award to MRW, and National Science Foundation grants IOS-1827645 and DEB-1856180 to ALS.

## Conflict of Interest

The authors declare no conflict of interests.

---

**Table 1: The *M. sookensis* genome can be split into two distinct subgenomes based on similarity to *M. nasutus* SF.** SD: Standard deviation.

**Figure S1: Range map of species used in this study.** Hand-drawn **r**anges are based on both sampled locations as well as observations and herbarium records. *M. sookensis* and *M. decorus* can be easily mis-identified as *M. nasutus* and *M. guttatus* respectively so ranges are an approximation.

**Table S1: Population sample information of all lines used in this study.** Depth for each line was calculated when mapped to the *M. guttatus* hardmasked v5 genome or to the *M. sookensis* FAN v1 genome.

**Table S2: Nucleotide diversity and divergence of focal species.** *Dxy* and *π* between *M. sookensis* subgenomes (A and B)*, M. nasutus* (N), and *M. guttatus* (G) calculated on fourfold degenerate sites.

**Figure S2 – *M. sookensis* contains 28 primary scaffolds**. Heatmap generated from HiC data and used to scaffold *M. sookensis* contigs. Scaffold identities are listed below x-axis.

**Figure S3: Homology between *M. sookensis* and its two progenitors. A.** *M. sookensis* v1.0 reference genome mapped to the *M. guttatus* v3 IM62 reference genome. Each *M. guttatus* chromosome shows homology with 2 *M. sookensis* chromosomes. Each dot indicates 75 Kb of homology. **B**. *M. sookensis* v1reference mapped against the *M. nasutus* v1 reference. Each *M. nasutus* chromosome shows homology with two *M. sookensis* chromosomes. Scaffolds are ordered from largest to smallest in size.

**Figure S4: M. sookensis subgenomes show high homology between subgenomes.** Circos plot generated from regions of homology between subgenomes A and B. Single ribbons represent 10 Kb regions of homology with a mapping quality of 60.

**Figure S5: NJ tree generated from 14,000 SNPs, bootstrapped 1000 times.**

**Figure S6: Maximum likelihood tree of *M. sookensis* subgenomes and diploid relatives in the *M. guttatus* species complex.** The ML tree was generated in IQ Tree using the TVM + F + R2 model from 1,711,738 parsimony informative sites with *M. dentilobus* as an outgroup.

**Figure S7: TWISST topology weights for gene trees with three “species”: *M. decorus*, northern *M. guttatus*, and southern *M. guttatus* + *M. nasutus*.** Data are from 16,754 gene trees rooted by *M. dentilobus* (not shown).

**Figure S8: All 15 simplified TWISST topology classes.** Letters correspond to Figure 4.

**Figure S9: Distribution of top three simplified TWISST topology classes across the 14 chromosomes.** Topologies were summed across windows of nine genes and plotted across the genome. Topology A is shown in green, Topology B in light blue, and topology C in navy.

**Figure S10: Distribution of the fourth most common TWISST topology class across the 14 chromosomes.** Topology D, which places *M. sookensis* subgenome B sister to *M. nasutus* + subgenome A could indicate regions of homoeologous recombination or gene conversion between subgenomes. Weights of topology D were summed across windows of 9 genes. Weightings are indicative of proportion of topology D in comparison to all other topologies.

## Notes

### Competing Interest Statement

The authors have declared no competing interest.

