## Supplementary figures and images for "Patterns of genomic variation reveal a single evolutionary origin of the wild allotetraploid *Mimulus sookensis*"

### Figure S1

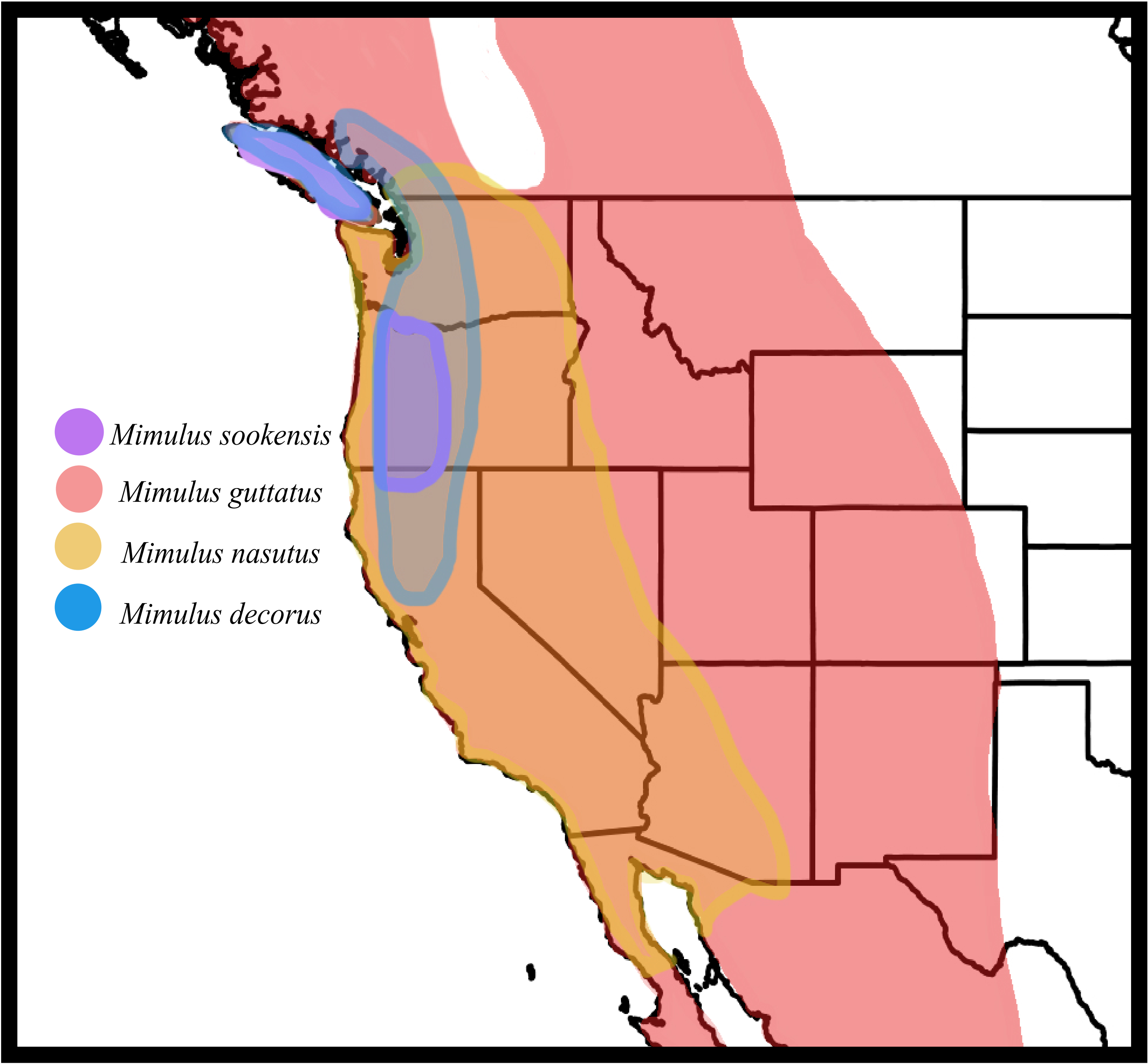

### Figure S2

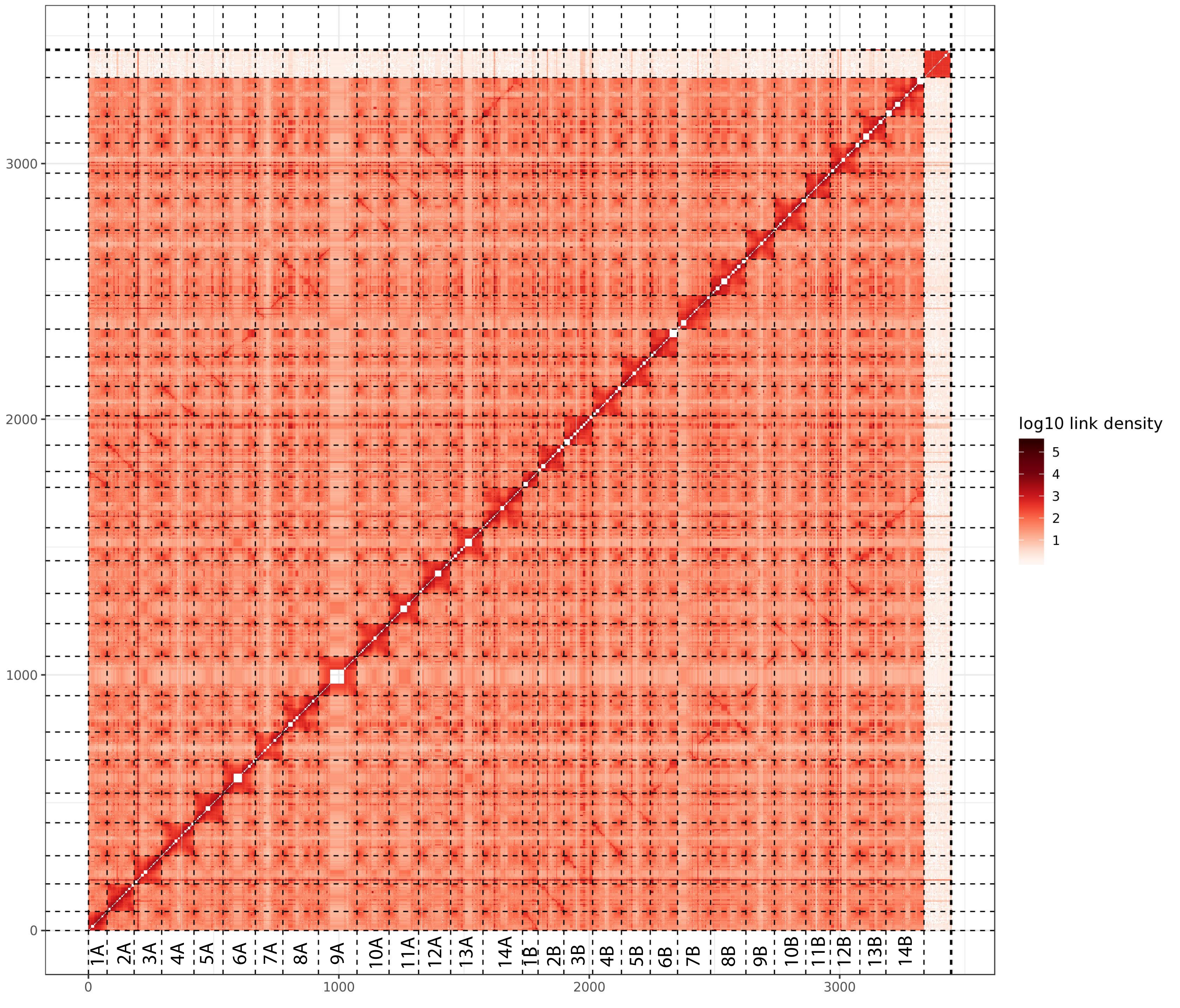

### Figure S3

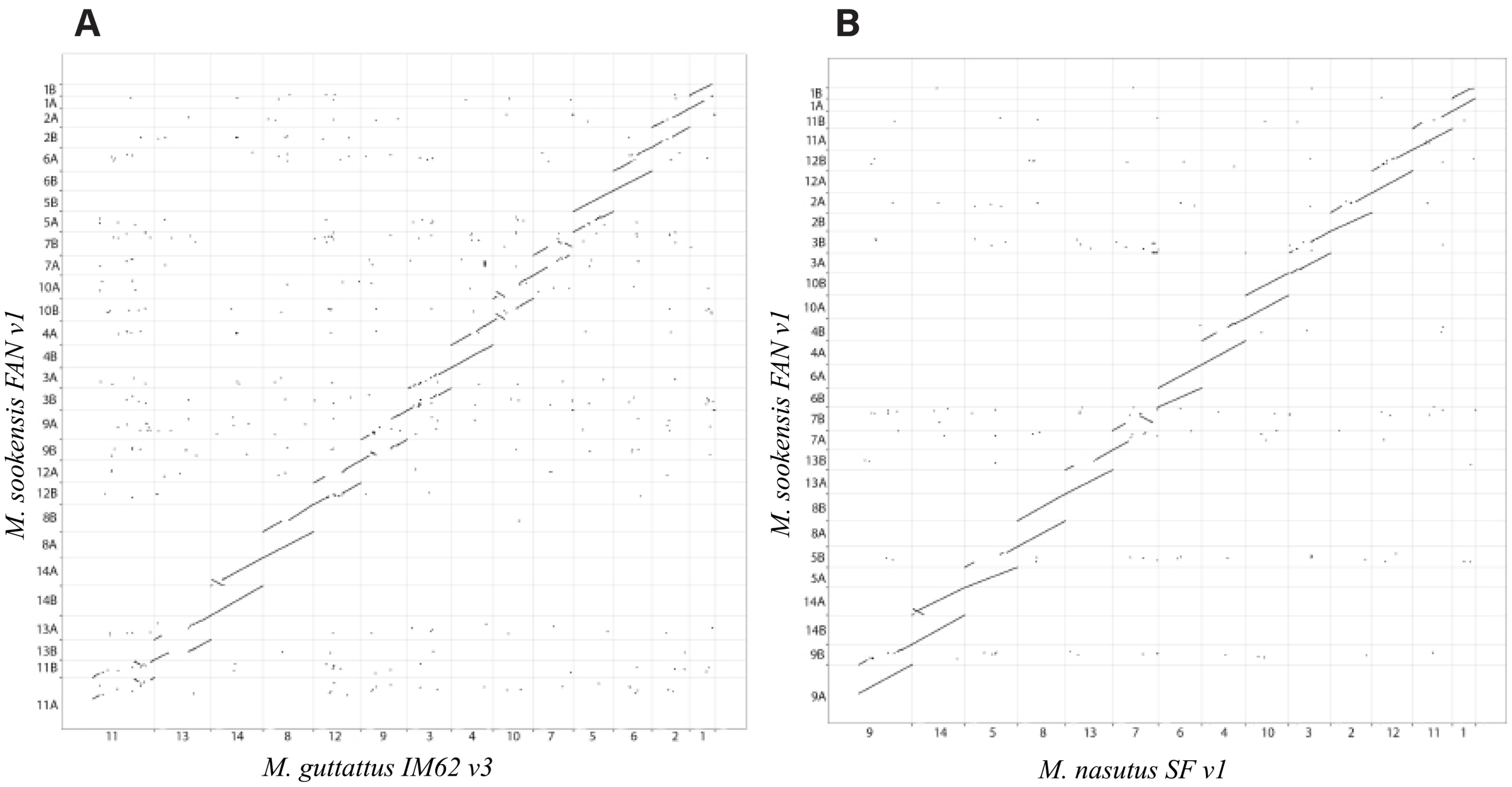

### Figure S4

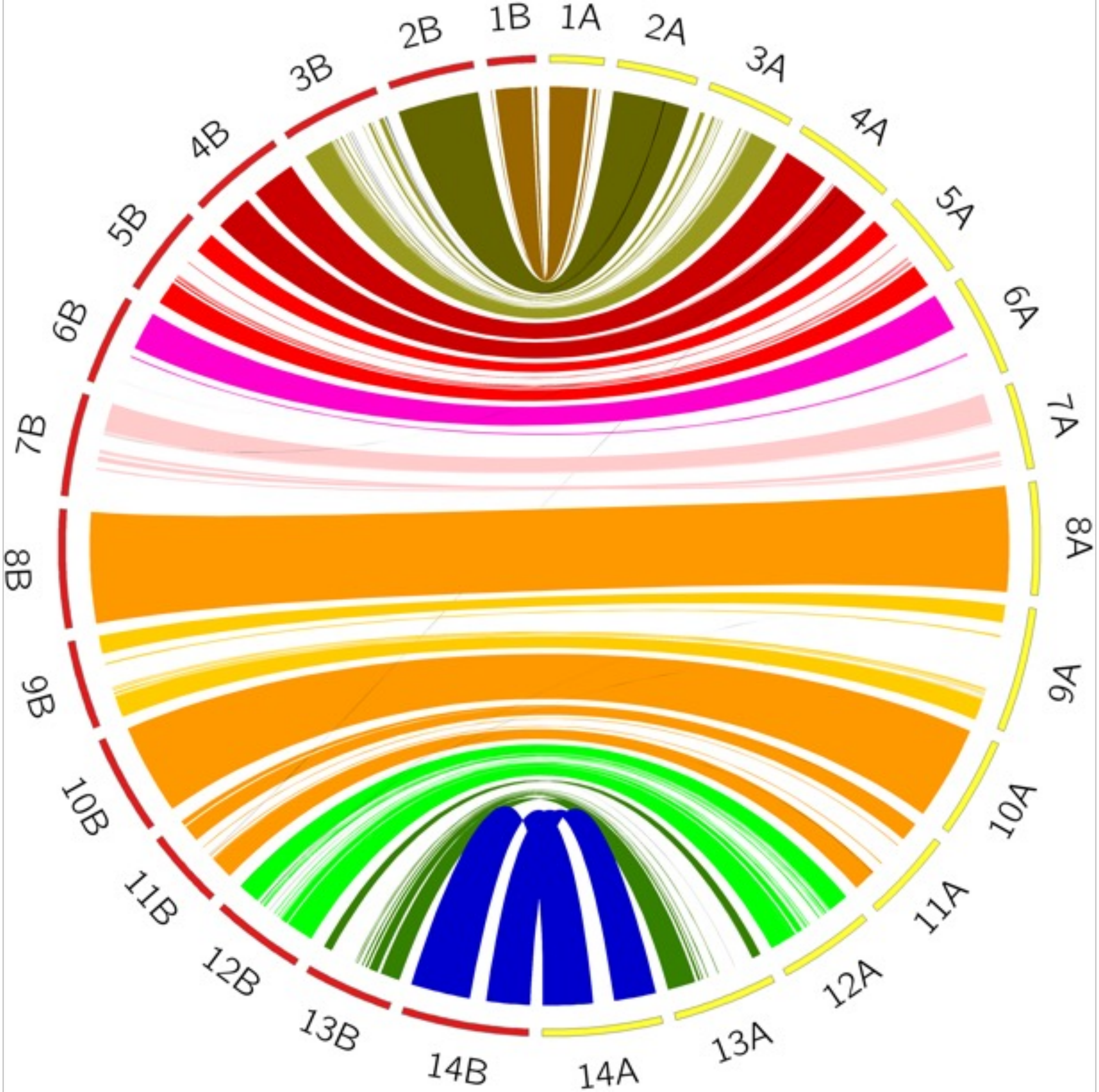

### Figure S5

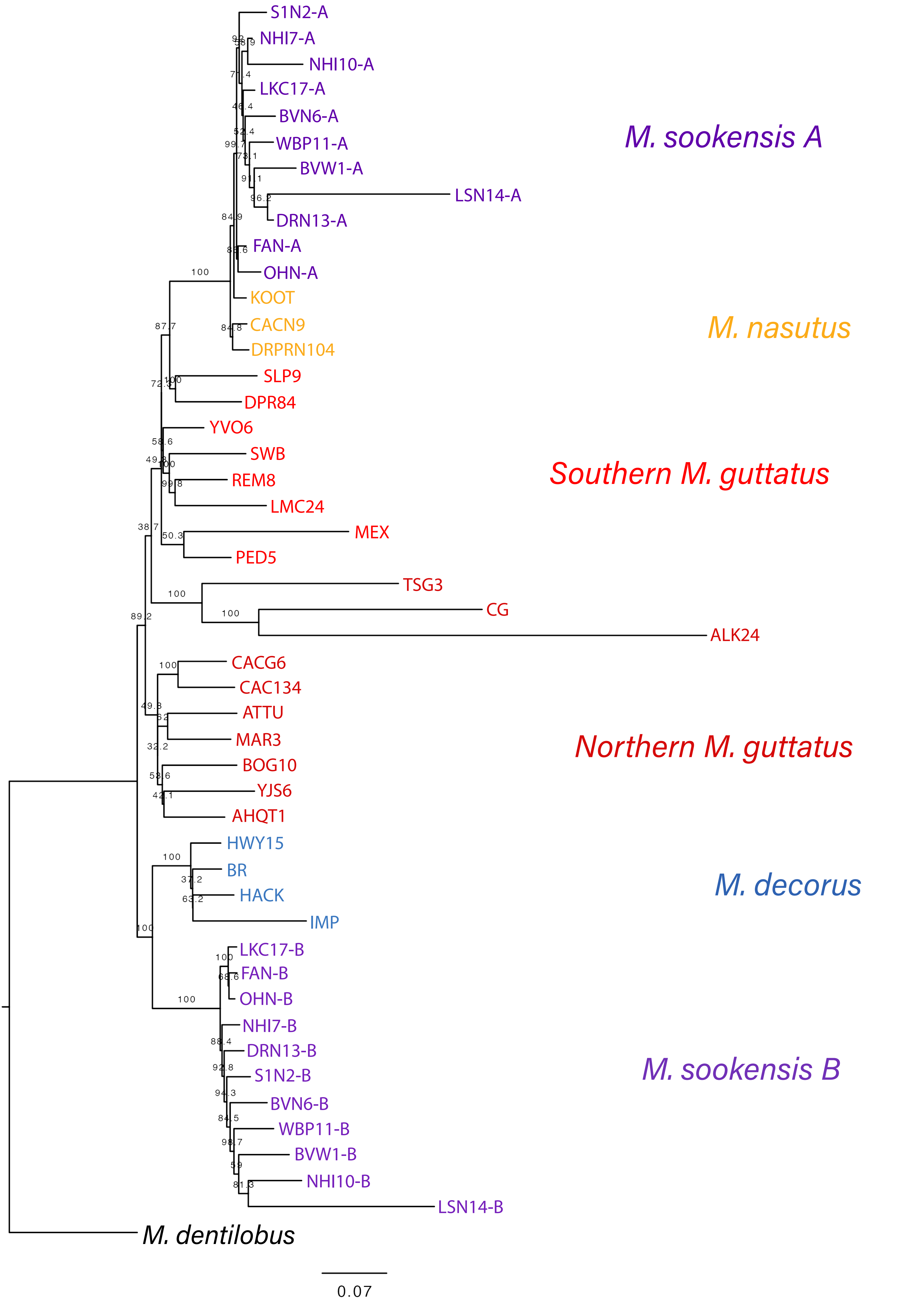

### Figure S6

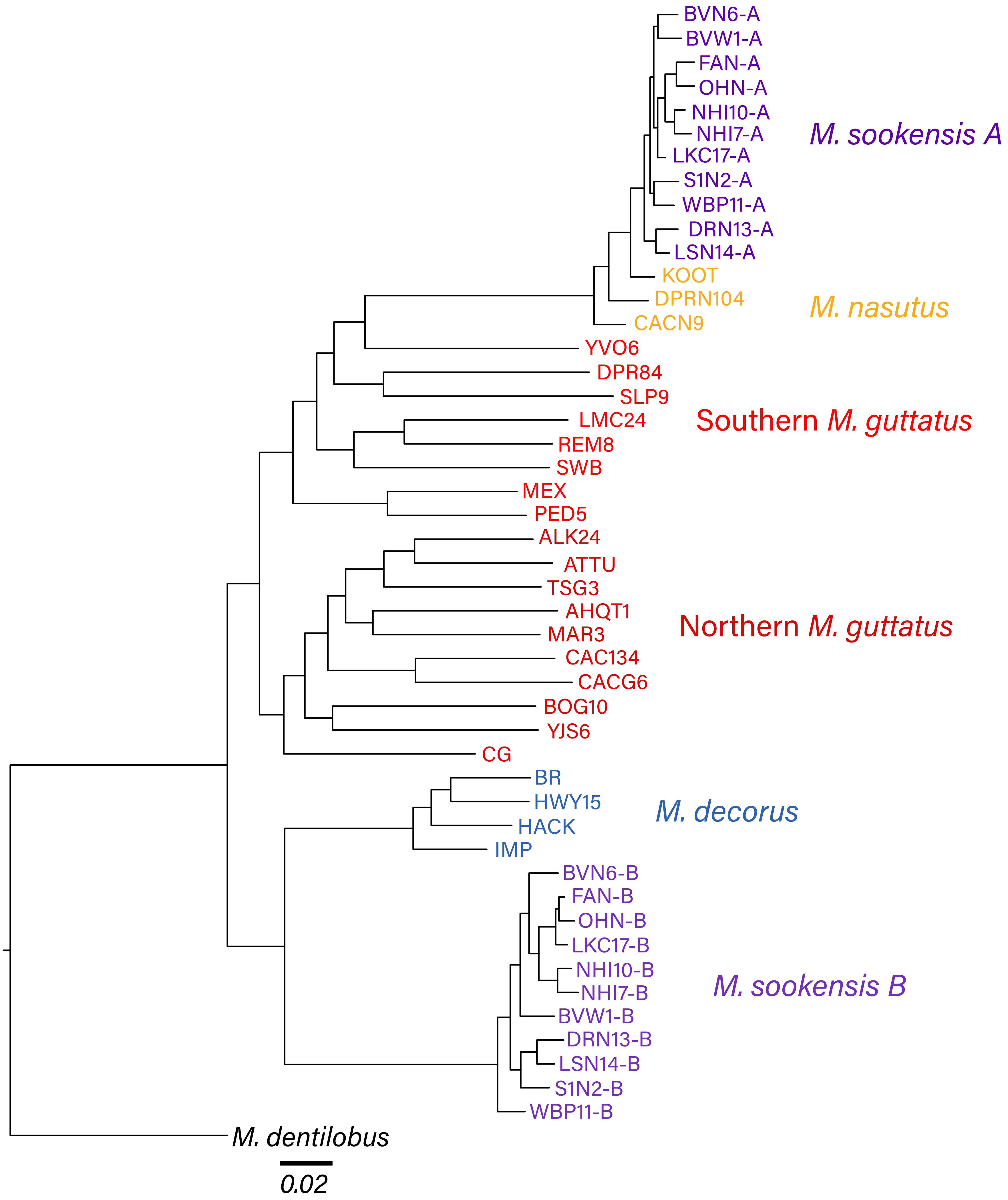

### Figure S7

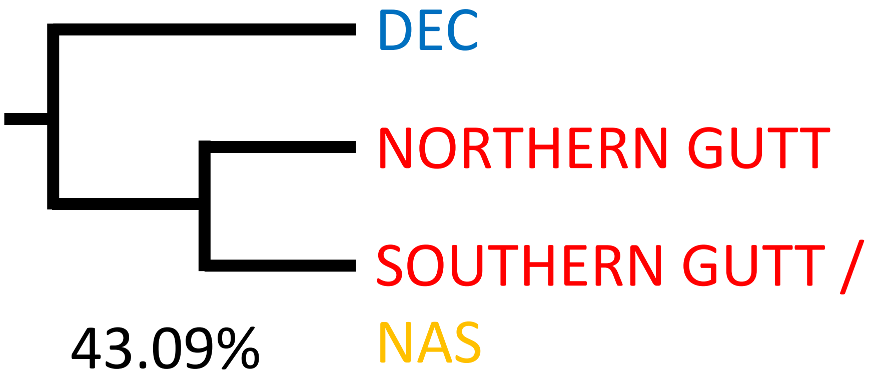

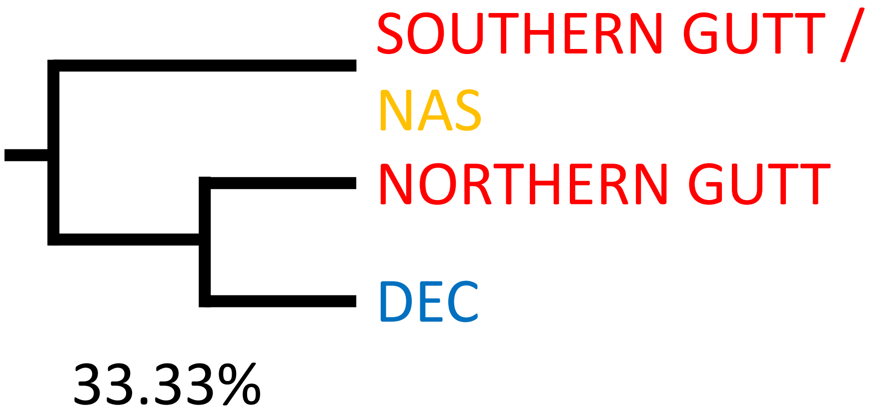

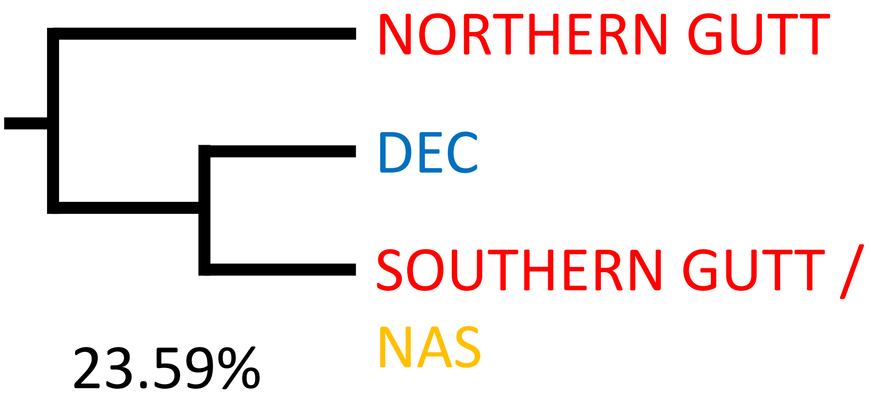

### Figure S9

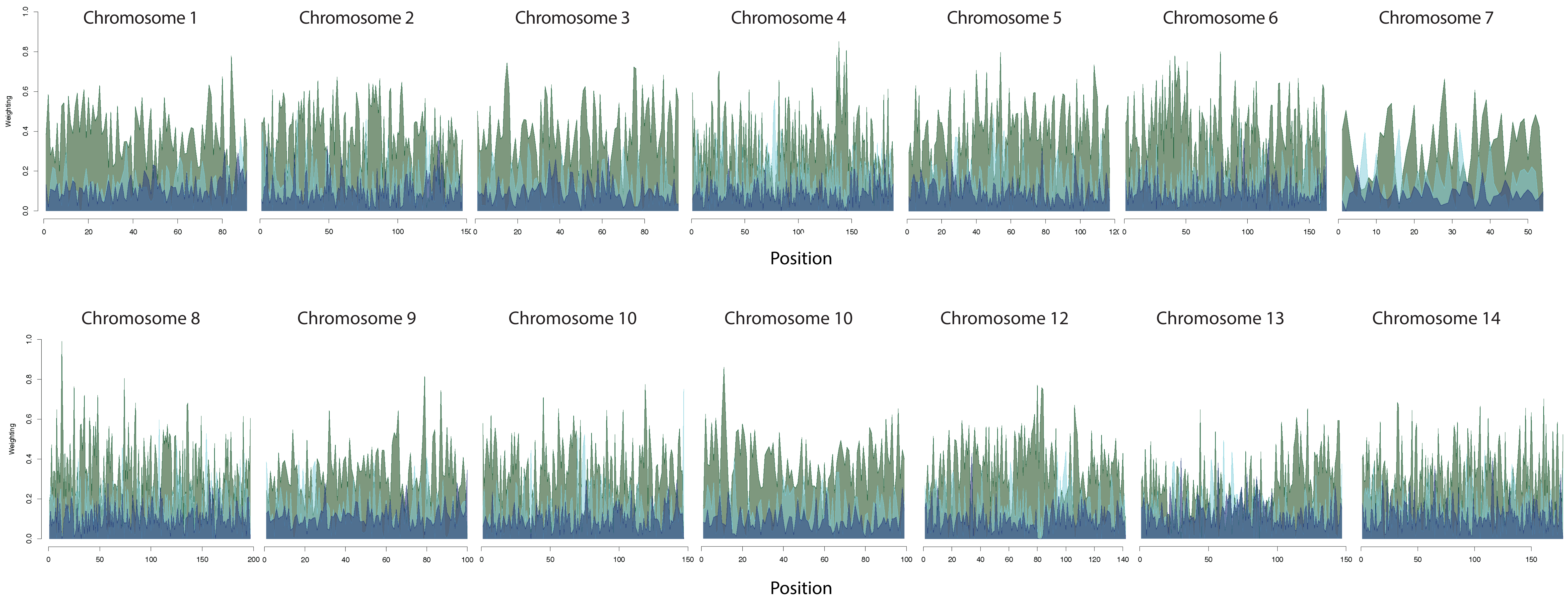

### Figure S10

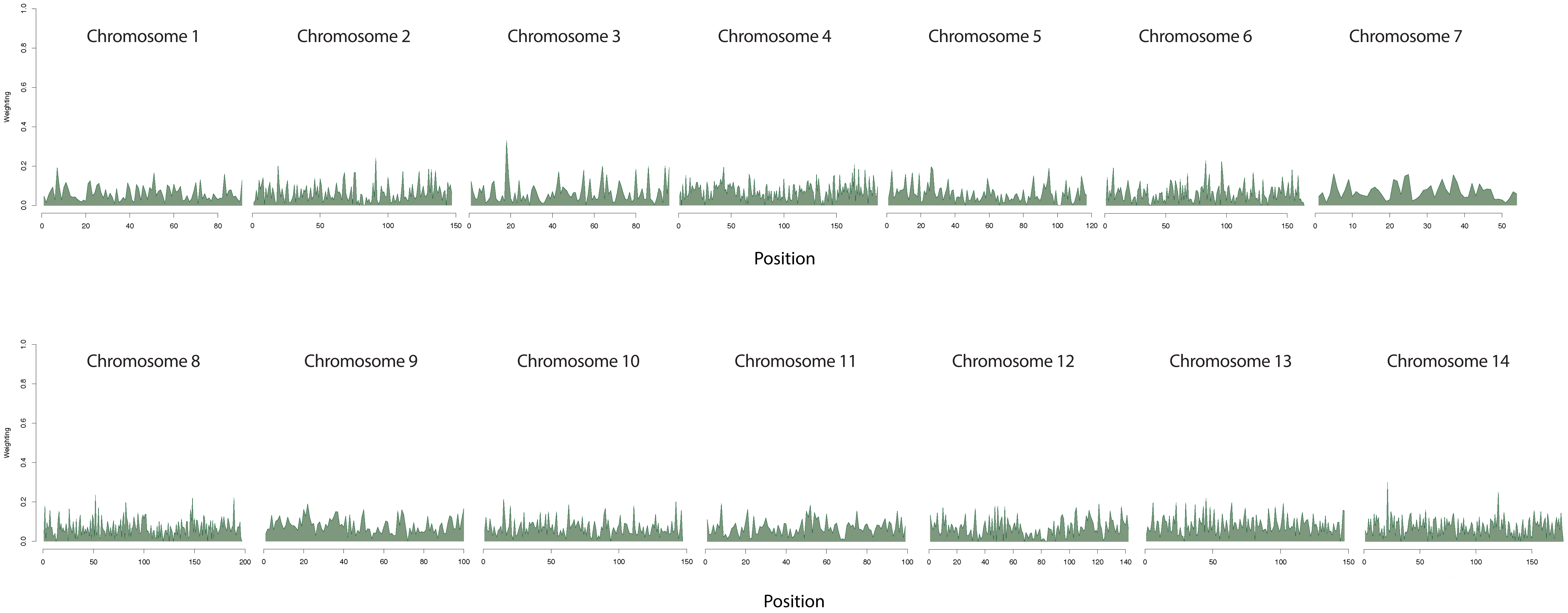
